# Comparison of CD4 T cell response in Plasmodium falciparum and vivax malaria

**DOI:** 10.1101/2025.04.28.651092

**Authors:** Mayimuna Nalubega, Megan SF Soon, Dean Andrew, Nicholas Dooley, Jessica R Loughland, Christian Engwerda, Enny Kenangalem, Ric N Price, Gabriela Minigo, Nicholas M Anstey, Damian A Oyong, Michelle J Boyle

## Abstract

**Background:** *Plasmodium falciparum* and *P. vivax* are parasites responsible for most malaria cases globally. In areas where these species co-exist, individuals gain protection from *P. vivax* more rapidly, and important biological differences between species may impact the immune response. CD4 T cells are key drivers of immunity to malaria, both as effector and helper cells, with T-follicular helper (Tfh) having key roles in antibody development. Comparative studies on CD4 T cell responses between these species are limited.

**Methods:** We assessed CD4 T cells in adults with either *P. falciparum* or *P. vivax* malaria. Activation and proliferation of CD4 T cells were measured *ex vivo*, and functional capacity determined by intracellular cytokine staining by flow cytometry.

**Results:** The phenotype, activation and proliferation of CD4 T cells and effector CD4 T cell subsets were comparable between species. However, within the peripheral (p)Tfh cell compartment, there was evidence for a skew towards pTfh1 cells in *P. falciparum*, and pTfh2 cells in *P. vivax*. Additionally, in *P. falciparum*, increased IL-10 production was detected, including within IL-21 producing CD4 T cells.

**Conclusion:** While activation and function of CD4 T cells in malaria are largely comparable, some species-dependent responses are detected within the pTfh cell compartment that may impact antibody development.

## Background

Malaria is a disease of global importance, with both *Plasmodium falciparum* and *P. vivax* causing significant disease morbidity and mortality[1]. While *P. falciparum* dominates in sub-Saharan Africa and is associated with the most severe morbidity and mortality, *P. vivax* has the widest geographical distribution and causes significant non-benign disease [1]. In areas where *P. falciparum* and *P. vivax* co-exist, individuals develop immunity faster to *P. vivax* irrespective of transmission intensity [2,3]. The rapid acquisition of immunity to *P. vivax* has been linked to a higher genetic diversity during infection compared to *P. falciparum* [4]. However, there are important biological differences between the two species including hypnozoite mediated relapses [5] and preferential life cycle of *P. vivax* in the spleen [6] which may also mediate significant differences in the immune responses to infection. Comparative studies in controlled human malaria models have suggested that species influence the cellular response to infection [7], however comparative studies in naturally acquired malaria are very limited.

CD4 T cells have important roles in protection from malaria in both naturally acquired and vaccine-induced immunity [8]. IFNɣ+ CD4 T cells (Th1) cells can drive parasite clearance through activation of macrophages [9] and have been associated with protection in both naturally acquired and controlled human *P. falciparum* malaria infection (CHMI) [10–15]. While data are limited in *P. vivax* malaria, the frequency of Th1 CD4 T cells negatively correlates with parasitemia in naturally acquired infection [16] suggesting malaria specific Th1 cells have important roles in immunity in both species. Additionally, CD4 T follicular helper (Tfh) cells, are essential to provide help to B cells to produce antibodies [17–19]. Activated peripheral (p)Tfh cells (CXCR5+PD1+) expand during both *P. falciparum* [19–21] and *P. vivax* infections in humans [16,22]. pTfh cells can be characterised into distinct subsets based on the expression of CXCR3 and CCR6 (pTfh1-CXCR3+CCR6–, pTfh17 - CXCR3–CCR6+, and pTfh2 - CXCR3–CCR6–), with different subsets associating with antibody development in a pathogen dependent manner [23]. In both *P. vivax* and *P. falciparum* malaria, pTfh responses are dominated by pTfh1 cells [16,19,21,22]. Aside from Th1 and pTfh cells, other CD4 T cells with roles in malaria immunity included regulatory subsets Type 1 regulatory T cells (Tr1) which co-produce IL-10 with IFNɣ, and dominate the malaria specific CD4 T cell immune response in children while are infrequent in immune adults [15,24–26]. FoxP3+ T regulatory cells (Tregs) have also been shown to expand during malaria, however in highly exposed children these cells are reduced with continuous exposure [27,28].

To date, few studies have specifically investigated species-associated differences in CD4 T cells during *P. falciparum* and *P. vivax* malaria. A recent study using *P. falciparum* and *P. vivax* blood stage CHMI in malaria naïve individuals highlighted potential important differences in the CD4 T cell immune response [7]. *P. vivax* induced a comparatively systemic higher type I inflammatory response, marked by elevated IFNɣ and CXCL9 levels, whereas *P. falciparum* induced higher T cell activation and terminal differentiation during acute infection. The CD38+ (Bcl2^lo^) activated memory and Treg CD4 T cells were higher in *P. falciparum*, and a higher Th1 transcriptional profile was detected compared to *P. vivax*. pTfh cells were not investigated in this study, and to the best of our knowledge no comparative studies of pTfh cells in the two parasite species have been published, nor have differences between the infections been comprehensively investigated in naturally acquired malaria. The aim of our study was to compare CD4 T cells activation and function in *P. falciparum* and *P. vivax* malaria in naturally acquired infection and specifically investigate pTfh cells which may have important roles in mediating faster acquisition of immunity to *P. vivax* by promoting antibody development.

## Results

### Activated CD4 T cells are comparable between P. falciparum and P. vivax malaria

To investigate species-dependent impacts of malaria on CD4 T cell activation we phenotyped responses in adults with acute *P. falciparum* (n=25) and *P. vivax* (n=12) malaria enrolled from previously conducted clinical studies using flow cytometry [29–31] (Sub-Cohort 1, **Supplementary Table S1**). CD4 T cell responses were also characterised in uninfected endemic controls of individuals from the same study site (EC, n=8). Within the CD4 T cell compartment, we identified Treg (FOXP3+ CD25+), peripheral Tfh (pTfh, CXCR5+ PD1+), and T helper cell subsets (Th1 – CXCR3+ CCR6-, Th17 – CXCR3-CCR6+, Th1-17 – CXCR3+ CCR6+), along with a chemokine negative subset which contained naïve CD4 T and Th2 cell responses (CXCR3-CCR6-) (**Supplementary Figure S1A**). Cell activation and proliferation status were determined by the expression of ICOS and Ki67, respectively (**Supplementary Figure S1A)**. Of total CD4 T cells, the frequency of ICOS+ and Ki67+ cells was significantly higher during both *P. falciparum* and *P. vivax* malaria compared to endemic controls, consistent with the expected cellular activation to acute infection (**Figure 1A**). Activation and proliferation were comparable between the two infecting species (**Figure 1A**), and neither ICOS+ nor Ki67+ was associated with parasitemia (**Supplementary Figure S1B**). At the CD4 T cell subset level, the proportions of pTfh cells and naïve/Th2 cells were higher in *P. falciparum* malaria compared to endemic controls with naive/Th2 also increased in *P. vivax*. Between *Plasmodium* species, there were no differences were observed across all CD4 T cell subset frequencies (**Figure 1B**), cell activation (**Figure 1C**) and proliferation (**Supplementary Figure S1B**).

**Figure 1:**
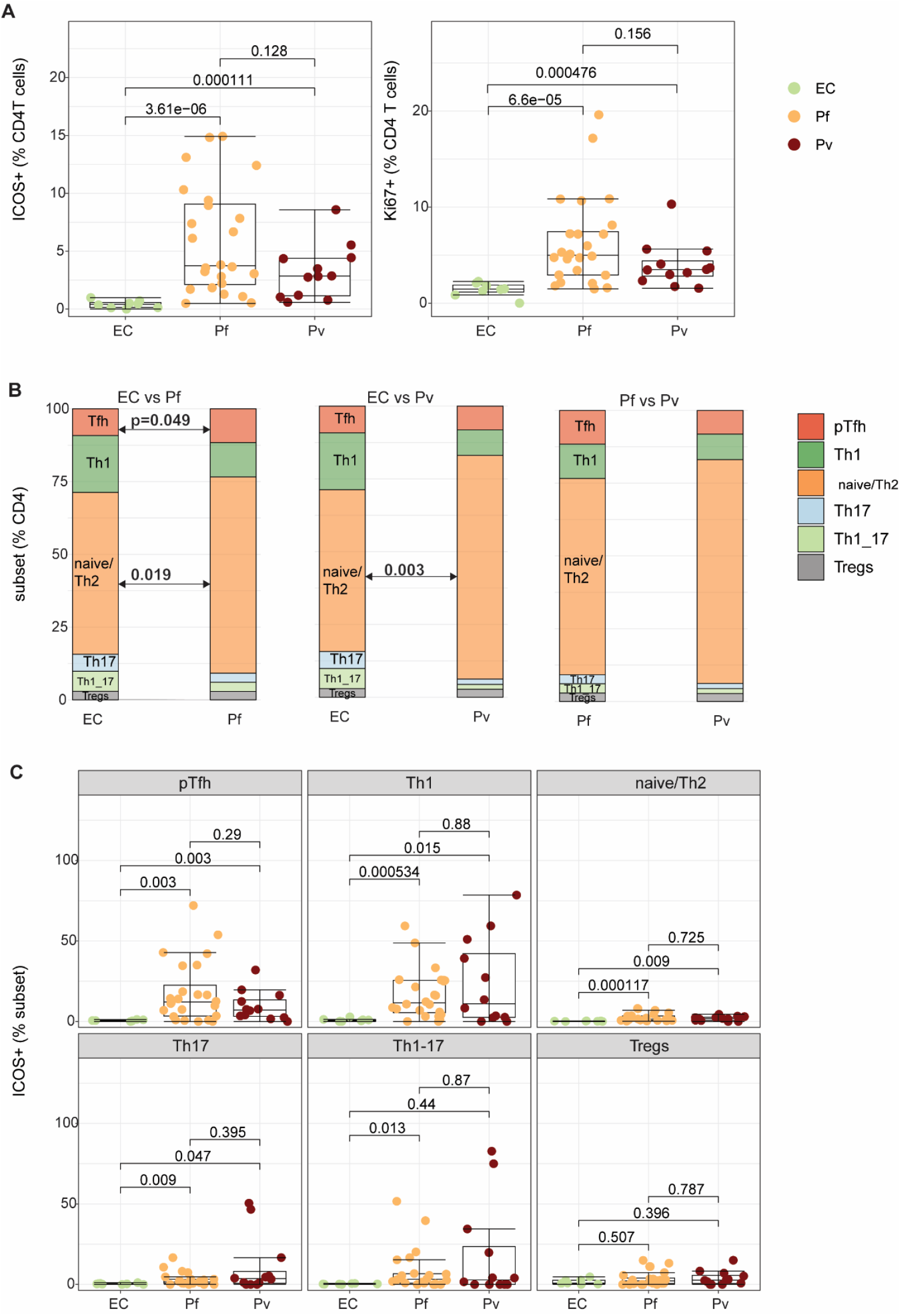
CD4 T cells are activated in acute P. falciparum and P. vivax malaria. CD4 T cell responses were assessed in adults with either P. falciparum (Pf, n = 25 or P. vivax (Pv, n = 12) malaria and in uninfected endemic controls (EC, n = 8). **A)** Frequency of ICOS+ (activation) and Ki67+ (proliferation) cells as a proportion of total CD4 T cells. **B)** Frequency of CD4 T cell subsets as a proportion of total CD4 T cells. **(C)** Frequency of ICOS+ cells as a proportion of each CD4 T cell subsets. For A and C box and whisker plots indicate first and third quartiles for the hinges, median line, and lowest and highest values no further than 1.5 interquartile range from the hinges for whisker lines. P-values indicate unpaired Wilcoxon signed-rank test.

### Activation of pTfh cell subsets is species-specific with a skew to pTfh1 in P. falciparum malaria

Since pTfh cells were activated and proportionally increased during malaria, we sought to determine if pTfh cell activation is species-specific at the subset level. Subsets of pTfh cells (CXCR5+PD1+) categorised into pTfh1 (CXCR3+CCR6-), pTfh2 (CXCR3-CCR6-), pTfh17 (CXCR3-CCR6+), and pTfh1-17 (CXCR3+ CCR6+) subsets, and activation and proliferation quantified (**Supplementary Figure S2**). The phenotypes of pTfh cell subsets were significantly different between *Plasmodium* species, with a higher proportion of pTfh1 cells in *P. falciparum* malaria and pTfh2 in *P. vivax* malaria when comparing between species (**Figure 2A**). The frequencies of ICOS+ pTfh1 and ICOS+ pTfh2 cells within the pTfh cell compartment increased in acute malaria compared to endemic controls regardless of species (**Figure 2B**). For proliferation, Ki67+ pTfh1 and pTfh2 cells were increased in *P. falciparum*, but only Ki67+ pTfh2 cells were increased in *P. vivax* (**Figure 2C)**. Further, significantly higher frequencies of ICOS+ and Ki67+ pTfh1 cells were observed in acute *P. falciparum* compared to *P. vivax* infection (**Figure 2B/C)**, together suggesting a pTfh1 skewed response in *P. falciparum* and relative pTfh2 skewed response in *P. vivax* malaria.

**Figure 2.**
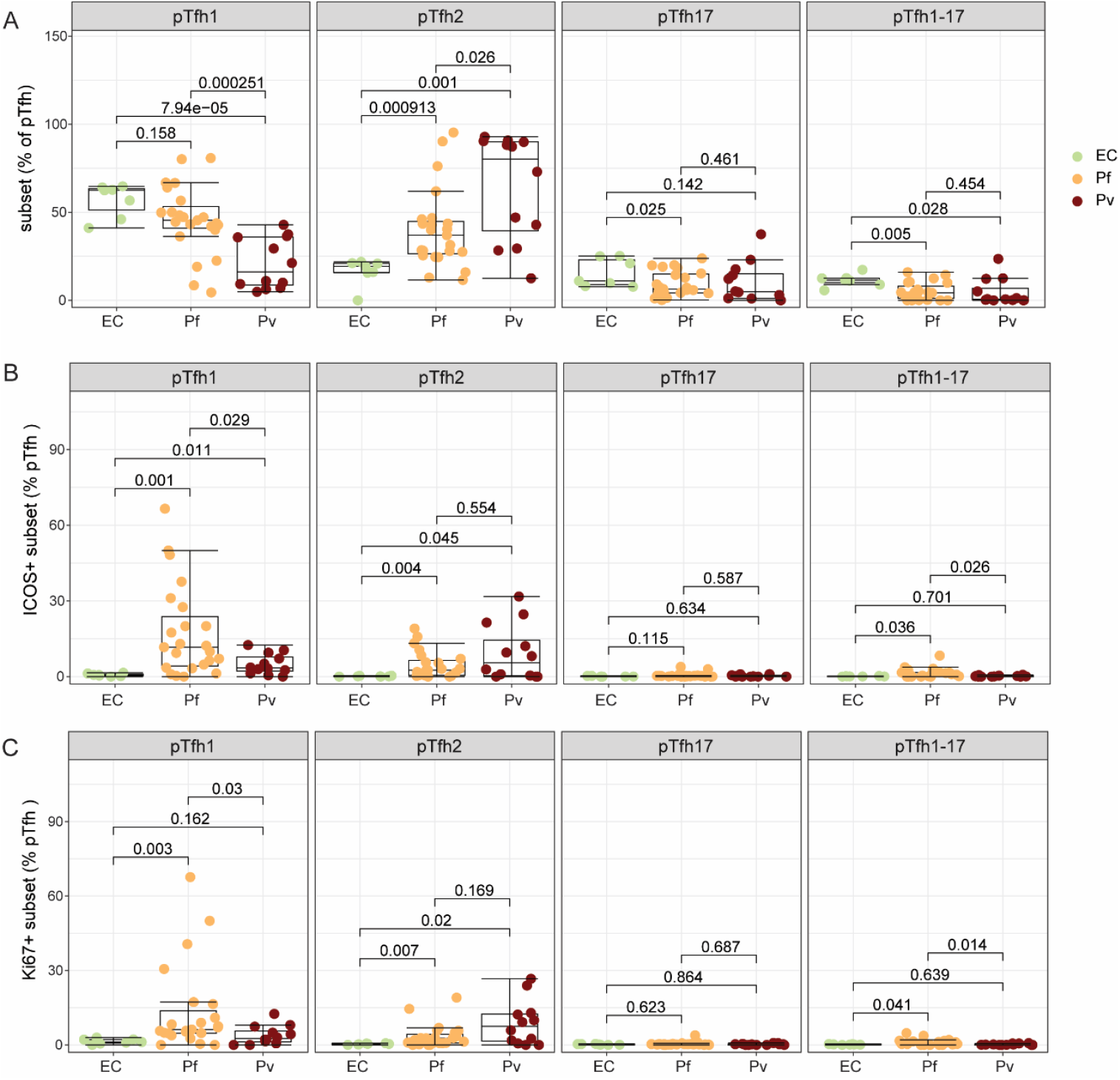
pTfh cell subset distribution and activation are species-specific during acute P. falciparum and P. vivax malaria. pTfh cell (CXCR5+PD1+ CD4 T cells) subsets were categorised into pTfh1 (CXCR3+ CCR6-), pTfh2 (CXCR3-CCR6-), pTfh17 (CXCR3-CCR6+) and pTfh1-17 (CXCR3+ CCR6+) cells. Responses were compared between uninfected endemic controls (EC, n = 8), and patients with either P. falciparum (Pf, n = 25), or P. vivax (Pv, n = 12) malaria. **A)** Frequency of pTfh cell subsets as a proportion of total pTfh cells. Frequency of **B)** ICOS+ and **C)** Ki67+ pTfh cells subsets as a proportion of total pTfh cells. Dots represent individual data coloured according to infection status groups. For all plots, box and whisker plots indicate first and third quartiles for the hinges, median line, and lowest and highest values no further than 1.5 interquartile range from the hinges for whisker lines. P-values indicate unpaired Wilcoxon signed-rank test.

### Unsupervised cell clustering analysis of CD4 T cells in P. falciparum and P. vivax malaria

To expand these findings, we designed a more comprehensive CD4 T cell panel to assess parasite species-dependent skewing on CD4 T cells and pTfh. Cellular responses were characterised in additional *P. falciparum* and *P. vivax* patients and uninfected endemic controls from the same clinical studies, along with malaria naïve Australian adults (Naïve) (Sub-Cohort 2, **Supplementary Table S2**). Non-naive CD4 T cells were identified based on CCR7 and CD45RA expression and further analysed with unsupervised cell clustering (**Figure 3A/B, Supplementary Figure 32A-C)**. Using this approach, we identified seven clusters which were annotated as pTfh (CXCR5+), regulatory T cells (Tregs) (CD25+ FOXP3+), Temra (CD45RA+), Th1 (CXCR3+ CCR6-CCR4-), Th2 (CXCR3-CCR6-CCR4+), Th17 (CXCR3-CCR6+ CCR4+), and Th1-17 (CXCR3+CCR6+) cells (**Figure 3A-B, Supplementary Figure S3A-C**). The proportions of CD4 T cell subsets were largely comparable regardless of malarial infection or naïve status, with the only significant differences observed being an increased proportion of pTfh cells in endemic controls compared to naïve individuals and in *P. falciparum* malaria compared to endemic controls (**Supplementary Figure S3D**). Consistent with the previous cohort (Sub-Cohort 1), the frequencies of ICOS+, CD38+, and Ki67+ cells across pTfh (p<0.05 for CD38+ and Ki67+) and Th1 (p<0.05 for ICOS+) cells were higher during acute *P. falciparum* and *P. vivax* malaria compared to endemic controls (**Figure 3C**). No significant differences were observed on activation and proliferation status within CD4 T cell subsets between *P. falciparum* and *P. vivax*, except for increased ICOS+ cells of pTfh cell subset in *P. vivax* compared to *P. falciparum* malaria (**Figure 3C, Supplementary Figure S4 A-C**).

**Figure 3.**
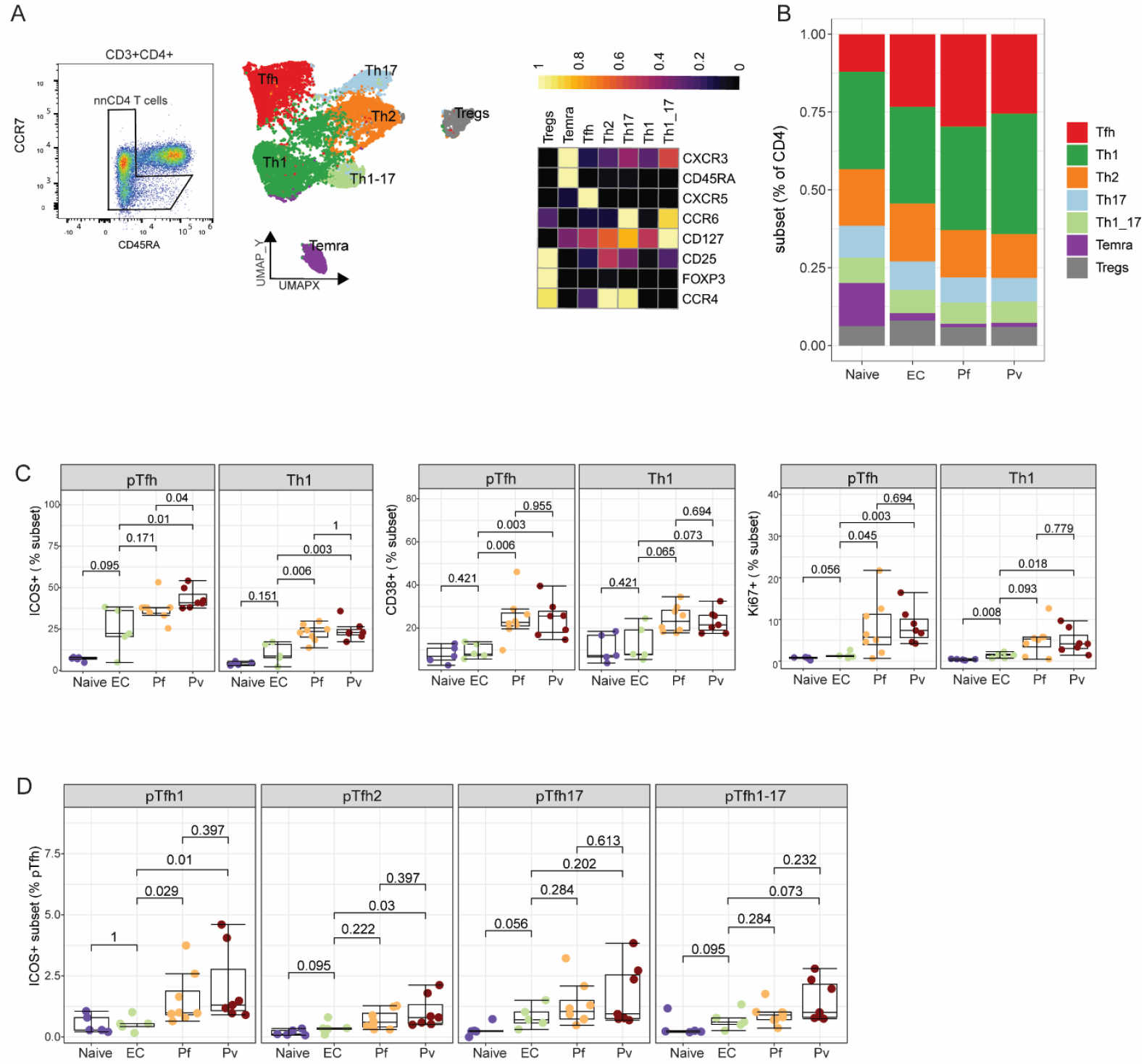
Unsupervised cell clustering analysis on CD4 T cell and pTfh cell subsets. CD4 T cells were analysed with a comprehensive panel and unbiased approaches in healthy malaria naïve Australians, healthy malaria uninfected controls, P. falciparum and P. vivax patients (Naive, n=5; EC, n=5; P. falciparum (Pf), n = 8; P. vivax (Pv), n = 7). **A)** Non-naïve CD4+ T cells were gated based on CCR7 and CD45RA expression. Non-naïve CD4 T cells were analysed with unbiased clustering and visualised with UMAP. Heatmap of normalised median expression of protein markers within annotated CD4 T cell subsets. **B)** Frequency of CD4 T cell subsets as proportions of non-naïve CD4 T cells. **C)** Frequency of activated ICOS+, CD38+ and Ki67+ pTfh and Th1 cells as proportions of CD4 T cells. **D)** Frequency of proliferating ICOS+ pTfh subsets as proportions of total pTfh. Data are individual responses coloured according to the control and malaria groups. Box and whisker plots indicate first and third quartiles for the hinges, median line, and lowest and highest values no further than 1.5 interquartile range from the hinges for whisker lines. P-values are Wilcoxon signed-rank test.

We further used unsupervised cell clustering analysis to categorise pTfh cell subsets based on CXCR3 and CCR6 expression (**Supplementary Figure S5A**). Consistent with the observation in Sub cohort 1, malaria infection induced activation of pTfh cells, and the frequencies of ICOS+, CD38+ and Ki67+ pTfh1 were higher in malaria infected compared to uninfected individuals (**Figure 3D, Supplementary Figure S5 C-D**). However, activation of pTfh2 cells was only detected in *P. vivax* malaria, with an increased frequency of ICOS+ pTfh2 cells compared to uninfected individuals (**Figure 3D)**. Furthermore, the proportion of ICOS+ cells within pTfh2 cells was higher in *P. vivax* than *P. falciparum* malaria (**Supplementary Figure S5E**). Taken together, our unbiased clustering analysis of CD4 T and pTfh cells during malaria are consistent with largely comparable CD4 T cell subset activation between species, and a skew towards pTfh1 cells in *P. falciparum* and a relative increase towards pTfh2 cells in *P. vivax* malaria.

### Increased IL-21+ CD4 T cells co-secreting IFNɣ and IL-10 in P. falciparum malaria

Finally, to elucidate functional potential of CD4 T cells in *P. falciparum* and *P. vivax* malaria, we stimulated cells from *P. falciparum* and *P. vivax* malaria patients and uninfected endemic and malaria naïve individuals with PMA/Ionomycin and quantified CD4 T cell production of IFNɣ, TNFa, IL-4, IL-17, IL-21 and IL-10, and degranulation marker CD107a (Sub-Cohort 2, **Supplementary Figure S6A**). PMA/Ionomycin cell stimulation increased all quantified cytokines and degranulation marker CD107a across all sample groups (**Supplementary Figure S6B)**. Following stimulation, the expression of IFNɣ, TNFa, IL-4, IL-17, IL-21. were comparable between all groups. In contrast, IL10 producing cells were higher in *P. falciparum* compared to *P. vivax* malaria and CD107a expression was elevated in *P. falciparum* compared to endemic controls (**Figure 4A**).

**Figure 4.**
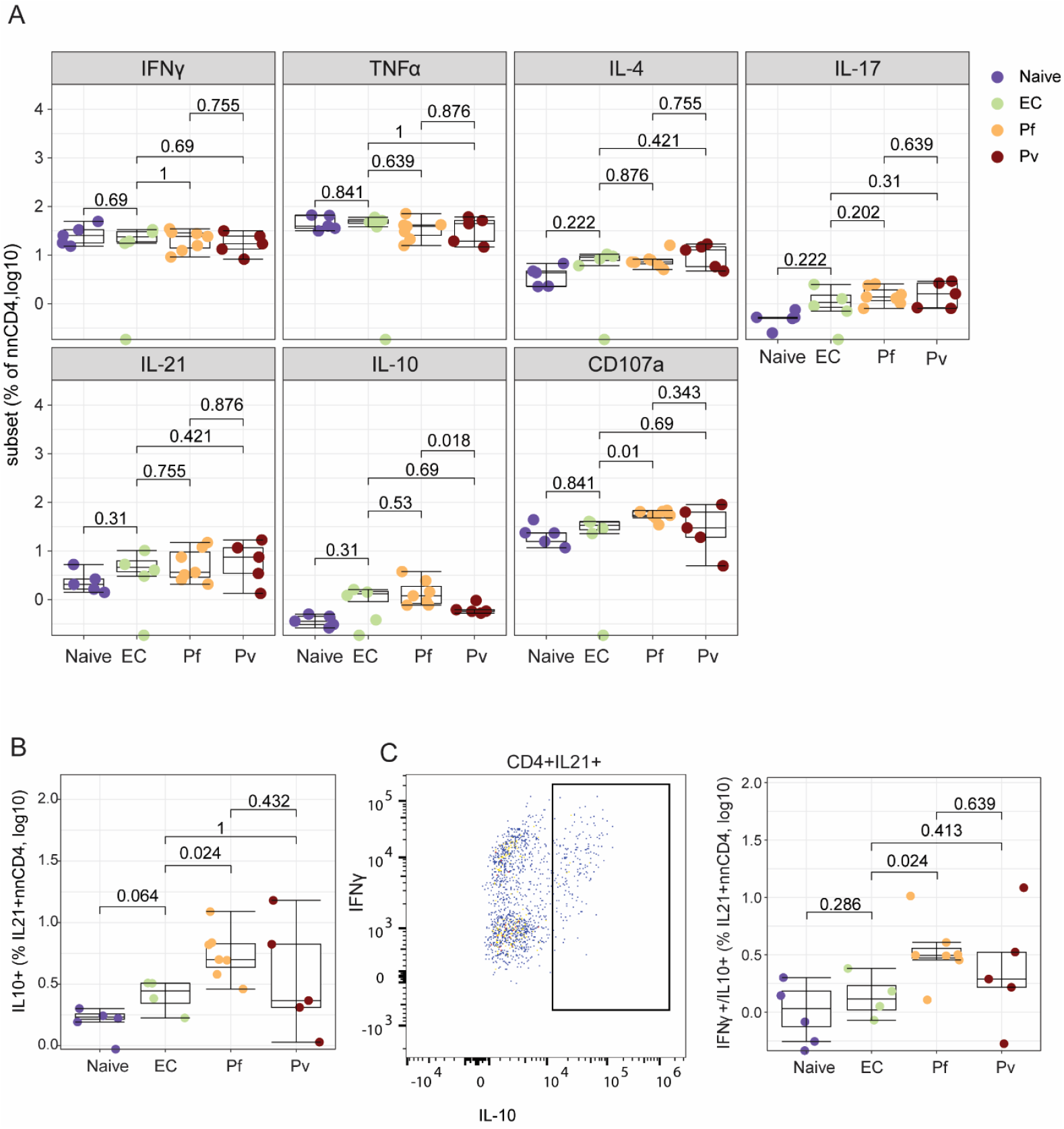
Functionality of pTfh subsets in Pf and Pv infections. PBMC samples were stimulated with PMA/Ionomycin to assess for cytokine secretion function between groups; healthy naïve Australians (Naïve, n = 5), endemic controls (EC, n = 5), P. falciparum (Pf, n = 7) and P. vivax (Pv, n = 5) malaria. **(A)** Frequency of cytokine producing cells in nnCD4 T cells. **(B)** IL-10+ producing cells within IL21+ nnCD4 T cells.**C)** Representative plot showing IL-10+ gating strategy and boxplots comparing frequency of IL-10+ and IFNy+ Dots represent individual data coloured according to the respective groups. Dots represent individual data coloured according to Tfh cell subsets. P-values were determined using paired Wilcoxon test. For all plots, box and whisker plots indicate first and third quartiles for the hinges, median line, and lowest and highest values no further than 1.5 interquartile range from the hinges for whisker lines.

IL-21 has key roles in antibody induction, modulating B cell responses through co-expression with other cytokines [23]. As such, we examined IL-21 co-production in CD4 T cells during malaria. Production of IL-10 within IL-21+ CD4 T cells was higher in *P. falciparum* malaria compared to endemic controls (**Figure 4B**). We found that these IL-21+ IL-10+ cells also expressed IFNɣ, and these cells were increased in *P. falciparum* malaria (**Figure 4C**). Co-production of other cytokines with IL-21 were similar across sample groups (**Supplementary Figure S7)**. Taken together, data suggest that pTfh subset activation and functional potential is influenced by infecting parasite species in malaria.

## Discussion

In this study, we compared CD4 T cell responses between naturally acquired *P. falciparum* and *P. vivax* malaria. Utilising both supervised and unbiased analyses, we show that CD4 T cell activation and phenotypes are generally similar between *Plasmodium* species. However, data suggest that pTfh cell subset responses were species-dependent, with increased skewing towards pTfh1 cells in *P. falciparum* and towards pTfh2 cells in *P. vivax* malaria. Beyond phenotypic differences, we also observe increased IL-10 production within CD4 T cells, and increased IL+10 co-expression with IL-21 during *P. falciparum* malaria. To the best of our knowledge, this is the first study that directly assess species-specific response of pTfh cells in malaria and taken together, inform our understanding of the immune acquisition process to *P. falciparum* and *P. vivax* infection.

CD4 T cells have multiple roles in immunity to malaria, with production of IFNg by Th1 cells contributing to parasite killing [32] and Tfh cells having essential roles in antibody development [33]. A recent experimental human malaria study comparing *P. falciparum* and *P. vivax* responses showed higher CD4 T cell activation in *P. falciparum* infection with Th1-associated gene signatures [7]. In comparison, our data in naturally acquired malaria shows that the magnitude of CD4 T cell activation and CD4 T cell subset distribution during acute malaria was largely species independent. We confirm these similarities of CD4 T cell responses using both supervised and unbiased analyses in two separate sub-cohorts. Although history of prior exposures is unknown in our cohort, in all cases, the adult individuals are presenting with symptomatic malaria, suggesting low levels of immunity. Nevertheless, study design and analytical differences including sample collection timepoint relative to peak parasitemia, may explain differences in findings between our and prior studies in controlled human malaria infection models [7].

While global CD4 T cell responses were comparable between *P. falciparum* and *P. vivax* infection, we observe evidence of species-specific response in pTfh cells. Specific subsets of pTfh cells can generate varying capacity of B cell help that impact antibody induction and maintenance. These pTfh subsets are thought to be driven by cytokine skewing in a context or disease specific manner [34]. Previous studies in *P. falciparum* [34,35] and *P. vivax* [37] malaria showed that pTfh1 cells are predominantly activated during infection. In this direct comparative study, our data suggest that pTfh cells in *P. falciparum* malaria have a greater skew towards pTfh1 cells while pTfh cell response during *P. vivax* infection comparably balanced to include pTfh2 cell subsets. We have previously shown that the pTfh2 cell subset is associated with functional antibodies [34]. Further, evidence suggest that pTfh1 cells are not associated with antibody response in Malian children [37], thus increased pTfh2 cells in *P. vivax* may drive relatively more rapid acquisition of antibodies in comparison to *P. falciparum*. However, both pTfh1 and pTfh2 cells are increased in Ugandan children with highest levels of protective malaria antibodies [38], and we have recently identified additional diversity with functional relevance within pTfh cells during controlled human malaria infection [40]. Thus, further studies are required to dissect the capacity of antigen-specific Tfh subsets cells in providing help to B cells and induce antibodies during malaria, and whether this is modulated by different *Plasmodium* species.

Along with species-specific pTfh cell subset activation, we also observed increased frequencies of IL-10 and IL21 co-producing cells CD4 T cells in *P. falciparum* malaria. Regulatory Tr1 CD4 T cells that secrete IL-10, together with IFNɣ, have previously been identified as a dominate CD4 T cell response in highly exposed populations [24,25], and these cells emerge rapidly, even after a single parasitic infection [41]. Consistent with our findings, IL-10 is also co-produced in IL-21+ CD4 T cells in *P. falciparum* infected pregnant women [42]. Although further studies are required to identify the underlying phenotypes and functions of IL-10+IL-21+ producing CD4 T cells and whether these cells are indeed a Tr1-like Tfh subset in human malaria, IL-10 producing Tfh cells have been found in chronic infections in mice models and were shown to support antibody production in the presence of persistent antigen[43]. Further in malaria murine models, IL-10 signalling was shown to promote GC B cell class switching and antibody responses [44], suggesting multifaceted functions of Tr1 cells than just being immunoregulatory. Given the importance of antibody development for both naturally acquired and vaccine induced protection, the role of IL-10 producing Tfh cells in human malaria infection is a focus of future research.

*Study limitations*. Our current study is limited by the small sample sizes across the two adult sub-cohorts with unknown prior malaria history from a single study site. Further analysis of CD4 T cell and Tfh cell activation in larger cohorts of different demographic profiles and age is required to further confirm and understand the broad relevance of our findings, particularly in children. Indeed, our previous studies have highlighted a role of age in Tfh cell activation and phenotypes in *P. falciparum* malaria [36,39]. Further, prior exposure to *P. falciparum* and *P. vivax* may impact Tfh cell activation however are unknown in this study. Additional experimental analysis is also required to dissect the impact of Tfh cell subset changes on antibody production and clinical outcomes, which requires longitudinal samples.

## Materials and methods

### Ethics statement

Written informed consent was obtained from all study participants. Studies were approved by the ethics committees of the Northern Territory Department of Health and Menzies School of Health Research (Darwin, Australia) #HREC 05-16, #HREC 03-64, #HREC 10-1397, QIMR Berghofer #HREC P3444 and #HREC P1479, Alfred Hospital Ethics Committee #188/23, the Indonesian National Institute of Health Research and Development (Jakarta, Indonesia) #NIHRD KS.02.01.2.1.4042, and the Oxford Tropical Research Committee (Oxford, United Kingdom) #OXTREC 013-04.

### Study participants

PBMC samples were obtained from previous clinical studies conducted in Timika between 2004 and 2007 [29–31]. Timika is a tropical forested lowland town in the Central-South Papuan province of Indonesia. This region has *P. falciparum and P. vivax* co-endemicity with unstable transmission. For the parent clinical studies, patients with blood smear-confirmed malaria and fever or a history of fever within the last 48 hours were enrolled in observational studies [29–31]. In a subset of trial participants, blood samples were collected at enrolment for PBMC isolation. For the current study, PBMC samples used were from adults with malaria infection categorised as mono-infection with *P. falciparum* or *P. vivax* by microscopy. Healthy endemic control adults from the same study site were also included and were malaria negative confirmed by blood smear, with no history of malaria infection in the previous month. Baseline clinical and laboratory details in each group are outlined in Suppl. Table 1,2. For the malaria naïve individuals, PBMCs were collected from healthy Australian adults with no history of prior malaria exposures.

### Flow cytometry

PBMCs were thawed in RPMI 1640 media (Gibco) containing L-Glutamine, 25nM HEPES,10% Fetal calf serum (FCS) and 0.02% benzonase nuclease. The cells were resuspended at a concentration of 1 x10^6^ cell/ 100µl media per well in 96 well plates and rested for 2 hours at 37°C, 5% CO_2._ The cells were stained with anti-CCR7 (PerCpCy5.5) at 37C for 45 minutes. The cells were washed and stained for 15minutes at room temperature with surface antibodies in **Supplementary Table S3 and S42**. Intracellular cytokine staining was performed using the FoxP3 Fix/Perm staining (eBioscience) as per the manufacturers’ manual. The intracellular cytokine stain antibodies are shown in **Supplementary Table S3 and S42**.

For detection of cytokine production, 1 x10^6^ cell/ 100ul media per well in 96 well plates were rested for overnight at 37°C, 5% CO_2._ The cells were stimulated with PMA (25ng/mL) and Ionomycin (1µg/mL) for 6 hours. A mix of protein transport inhibitors Brefeldin A and monensin was added during the last four hours of stimulation. To maximise surface staining of CXCR3, CCR6, CXCR5 and CCR7 due to their downregulation after exposure to PMA and Ionomycin, anti-CXCR3 PE CF594, anti-CCR6 BV 605, anti-CCR7 PerCP-Cy5 and human Fc block (BD eBiomedicine) were added to the culture during stimulation. After stimulation, cells were stained for 15 minutes at room temperature with the live dead blue stain (InvitrogenTM) and surface labelled antibodies in **Supplementary Table 3**.

Intracellular staining was performed using the BD Cytofix/Cytoperm kit (BD Biosciences) according to manufacturer’s instructions. Details of the surface and intracellular cytokine antibodies are in Supplementary Table 3. All cells were resuspended in 2% FCS/PBS prior to acquisition with Beckman Coulter Gallios for Sub Cohort 1 data or using the AURORA spectral flow cytometer for Sub Cohort 2 data.

Data analysis was performed using FlowJo (version 10.7.2). Populations were gated as indicated; CD4 T cells (**Supplementary Figure S1**), cytokines (**Supplementary Figure S2**). Boolean “AND” gates were used for co-expression

### Unsupervised clustering and dimensional reduction

Cell subsets were identified using the unsupervised clustering algorithm FlowSOM [45] followed by dimensional reduction using UMAP [45].The Spectre package (v1.0) was used in R to perform clustering and dimensional reduction analyses [47]. Briefly, non-naive CD4 T cells were gated and exported using FlowJo prior to the clustering analysis. Over clustering approach (k=40) was used to generate abundant number of clusters that were then individually grouped and annotated based on the expression of relevant markers. Annotated cell clusters were visualised on UMAP plots at 10,000 total cells. Cell count and proportion data were calculated for downstream statistical analyses in R.

### Statistics

Group comparisons between naive, endemic controls (EC), *P. falciparum* and *P. vivax* were performed using unpaired non-parametric Wilcoxon test. All analyses were performed in R (version 4.1.3) and Spectre (version 1.1.0).

## Acknowledgements

Burnet Institute, QIMR-Berghofer and Menzies School of Health Research acknowledge the traditional custodians of the lands where they are located, the Boonwurrung people of the Kulin Nation, the Turrbal and Jagera people, and the Larrakia Nation. We thank staff of the local hospital, Indonesian Ministry of Health, and everyone involved in the Timika studies.

This work was supported by the National Health and Medical Research Council of Australia (Project Grant 1125656 and Career Development Fellowship 1141632 to MJB, Senior Principal Research Fellowship 1135820 to NMA), the CSL Centenary Fellowship to MJB, and the Snow Medical Foundation Fellowship 2022/SF167 to MJB, the Charles Darwin University (PhD scholarship to DAO), Menzies School of Health Research (PhD Top-Up Award to DAO), Channel 7 Children’s Research Foundation (grant to MJB and GM). The Burnet Institute is supported by the National Health and Medical Research Council for Independent Research Infrastructure Support Scheme and the Victorian State Government Operational Infrastructure Support.

## Author contributions

MN, MSFS, DA, ND, JRL, DAO generated data, supervised by MJB, GM, RP, NMA

MN and DAO analysed data, supervised by MJB

MJB, GM and DAO conceptualised study

EK, RP, GM, NMA provided clinical samples

MN, DAO and MJB led manuscript writing with contribution and approval from all authors

## Data availability statement

All data required for conclusions is available in manuscript. Additional data is available from corresponding authors upon reasonable request and appropriate agreements.

