## Supplementary Figures and Tables for "Comparison of CD4 T cell response in Plasmodium falciparum and vivax malaria"

**Supplementary Tables and Figures**

**Supplementary Tables**

**Supplementary Table S1: Sub cohort 1: Demographics and clinical data**

| Characteristics | Endemic healthy controls (EC), N = 8 | *P. falciparum* (Pf), N = 25 | *P. vivax* (Pv), N = 12 |
|---|---|---|---|
| Male (N, %) | 7 (88%) | 12 (48%) | 6 (50%) |
| Age Median (IQR) | 29 (27, 31) | 22 (20, 28) | 25 (21, 30) |
| Hb (g/dL) | 13.25 (12.25, 14.90) | 10.60 (9.40, 12.20) | 11.80 (10.05, 12.20) |
| Parasites (per/µL) # | NA | 14,040 (6,836, 48,000) | 6,895 (4,915, 9,630) |

*Missing gender information (N; Pf = 2, Pv = 1)
#Blood film diagnosis

**Supplementary Table S2: Sub cohort 2 demographics and clinical data**

| Characteristics | Malaria naive (Naive), N=5 | Endemic healthy controls (EC), N = 5 | *P. falciparum* (Pf), N = 8 | *P. vivax* (Pv), N = 7 |
|---|---|---|---|---|
| Male (N, %) | 2(40%) | 3 (75%) | 4 (50%) | 3(43%) |
| Age, Median (IQR) | 36 (30,43) | 26 (21, 30) | 26 (23, 27) | 25 (20, 28) |
| Hb (g/dL) | NA | 12.20 (10.68, 14.18) | 11.90 (11.30, 12.60) | 11.40 (10.00, 12.23) |
| Parasites (per/µL) # | NA | NA | 186,648 (7,253, 275,528) | 4,997 (4,589, 7,282) |

#Blood film diagnosis

#### Supplementary Tables and Figures

##### Supplementary Table 3: Antibodies for phenotyping of Tfh cells *exvivo* via flow cytometry

| Fluorophore | Marker | Clone | Catalogue number | Manufacturer | Dilution |
|---|---|---|---|---|---|
| BUV496 | CD8 | RPA-TS | 612942 | BD | 1/100 |
| BUV563 | CD45RA | H100 | 565702 | BD | 1/400 |
| BUV737 | CD25 | 2A3 | 612807 | BD | 1/200 |
| BUV805 | CD3 | SK7 | 612893 | BD | 1 in 50 |
| BV480 | CD38 | HIT2 | 566137 | BD | 1 in 50 |
| BV510 | CD14 | M5E2 | 301842 | BioLegend | 1 in 100 |
| BV510 | CD19 | H1B19 | 30184 | BioLegend | 1 in 100 |
| BV570 | CD127 | A019D5 | 351308 | BioLegend | 1 in 50 |
| BV605 | CCR4 | L291HL1 | 301842 | BioLegend | 1 in 50 |
| BV650 | CCR6 | 11A9 | 359418 | BioLegend | 1 in 50 |
| BV711 | CXCR5 | J252D4 | 563922 | BioLegend | 1 in 50 |
| BV785 | CD4 | OKT4 | 317442 | BioLegend | 1 in 50 |
| PE-CF594 | CXCR3 | IC6/CXCR3 | 302906 | BD | 1 in 50 |
| PE-Cy7 | PD-1 | EH12.1 | 562451 | BD | 1 in 50 |
| AF700 | CD161 | HP-3G10 | 563473 | BioLegend | 1 in 50 |
| APCCy7 | ICOS | C398.4A | 313530 | BioLegend | 1 in 25 |
| Live dead Blue | | | L23105 | Invitrogen | 1 in 5000 |
| **Stain at 37C,45minutes** | | | | | |
| PerCP-Cy5.5 | CCR7 | G043H7 | 353220 | BD | 1 in 12.5 |
| **Intracellular staining** | | | | | |
| BUV395 | Ki67 | B56 | 584071 | BD | 1 in 250 |
| PE | GZMB | GB11 | 561142 | BD | 1 in 400 |
| AF647 | FOXP3 | 320114 | 206D | BioLegend | 1 in 50 |

**Supplementary Tables and Figures**

**Supplementary Table 4: Antibodies for cytokine measurement after stimulation with PMA/Ionomycin**

| **Fluorophore** | **Marker** | **Clone** | **Catalogue number** | **Manufacturer** | **Dilution** |
|---|---|---|---|---|---|
| BUV496 | CD8 | RPA-TS | 612942 | BD | 1 in 100 |
| BUV563 | CD45RA | H100 | 565702 | BD | 1 in 400 |
| BUV805 | CD3 | SK7 | 612893 | BD | 1 in 50 |
| BV510 | CD14 | M5E2 | 301842 | BioLegend | 1 in 100 |
| BV510 | CD19 | H1B19 | 30184 | BioLegend | 1 in 100 |
| BV605 | CCR4 | L291HL1 | 301842 | BioLegend | 1 in 100 |
| BV650 | CCR6 | 11A9 | 359418 | BioLegend | 1 in 50 |
| BV711 | CXCR5 | J252D4 | 563922 | BioLegend | 1 in 50 |
| BV785 | CD4 | OKT4 | 317442 | BioLegend | 1 in 50 |
| PE-CF594 | CXCR3 | IC6/CXCR3 | 302906 | BD | 1 in 50 |
| PE-Cy7 | PD-1 | EH12.1 | 562451 | BD | 1 in 50 |
| APC-Fire | Vd2 | B6 | 331419 | BioLegend | 1 in 50 |
| **Stain at 37C,45minutes** | | | | | |
| PerCP-Cy5.5 | CCR7 | G043H7 | 353220 | BD | 1 in 25 |
| **Intracellular staining** | | | | | |
| BUV395 | IFNy | B27 | 563563 | BD | 1 in 50 |
| BV421 | IL-10 | JES3-9D7 | 564053 | BD | 1 in 100 |
| BV750 | TNFa | Mab11 | 566359 | BD | 1 in 100 |
| FITC | IL-17 a | BL168 | B252099 | BioLegend | 1 in 25 |
| PE | IL-21 | 3A3-N2.1 | 560463 | BD | 1 in10 |
| APC | IL-4 | MP4-25D2 | 500812 | BioLegend | 1 in 10 |

#### Supplementary Tables and Figures

#### Supplementary Figures

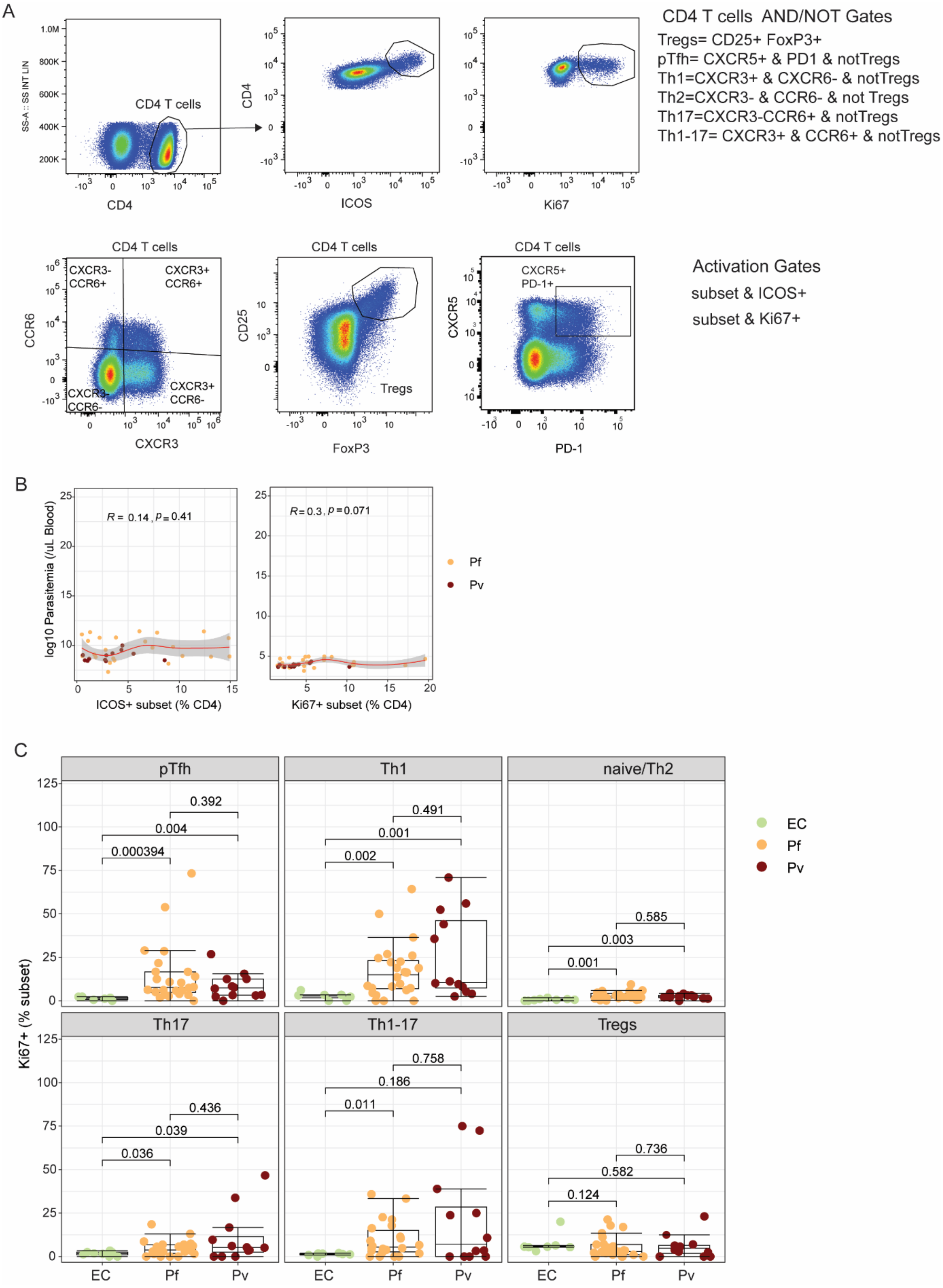

### Supplementary Tables and Figures

***Supplementary Figure S1: Gating and proliferation of CD4 T cells in P. falciparum and P. vivax malaria (Sub-Cohort 1)***

*CD4 T cell responses were assessed in adults with P. falciparum (Pf, n = 25) and P. vivax (Pv, n = 12) compared with endemic healthy controls (EC, n = 8).* ***A)*** *Gating strategy to identify CD4 T cell subsets and activation and proliferation.* ***B)*** *Correlation between ICOS+ and Ki67+ CD4 T cells and parasitemia. R and p are spearman's correlations.* ***C)*** *Activation (ICOS+) as a proportion of each CD4 T cell subsets in healthy controls and patients.*

**Supplementary Tables and Figures**

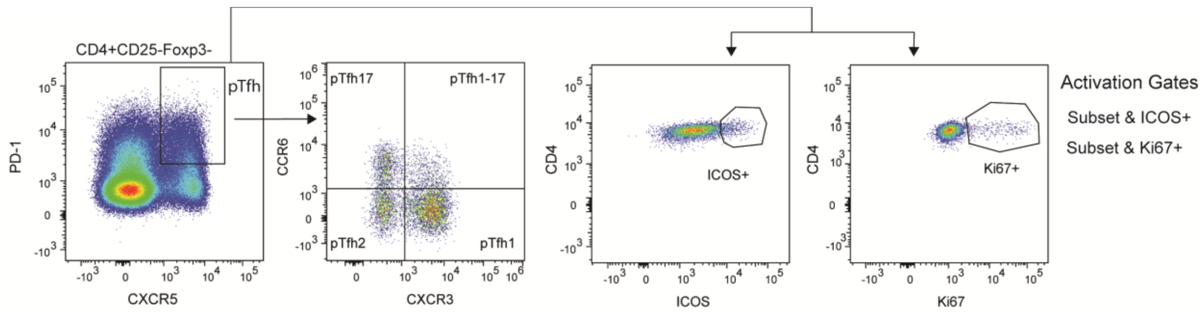


***Supplementary Figure S2: pTfh gating and subset activation (Sub-Cohort 1)***

*CD4 T cell responses were assessed in adults with P. falciparum (Pf, n = 25) and P. vivax (Pv, n = 12) compared with endemic healthy controls (EHC, n = 8). Gating strategy to identify pTfh cells and activation and proliferation.*

#### Supplementary Tables and Figures

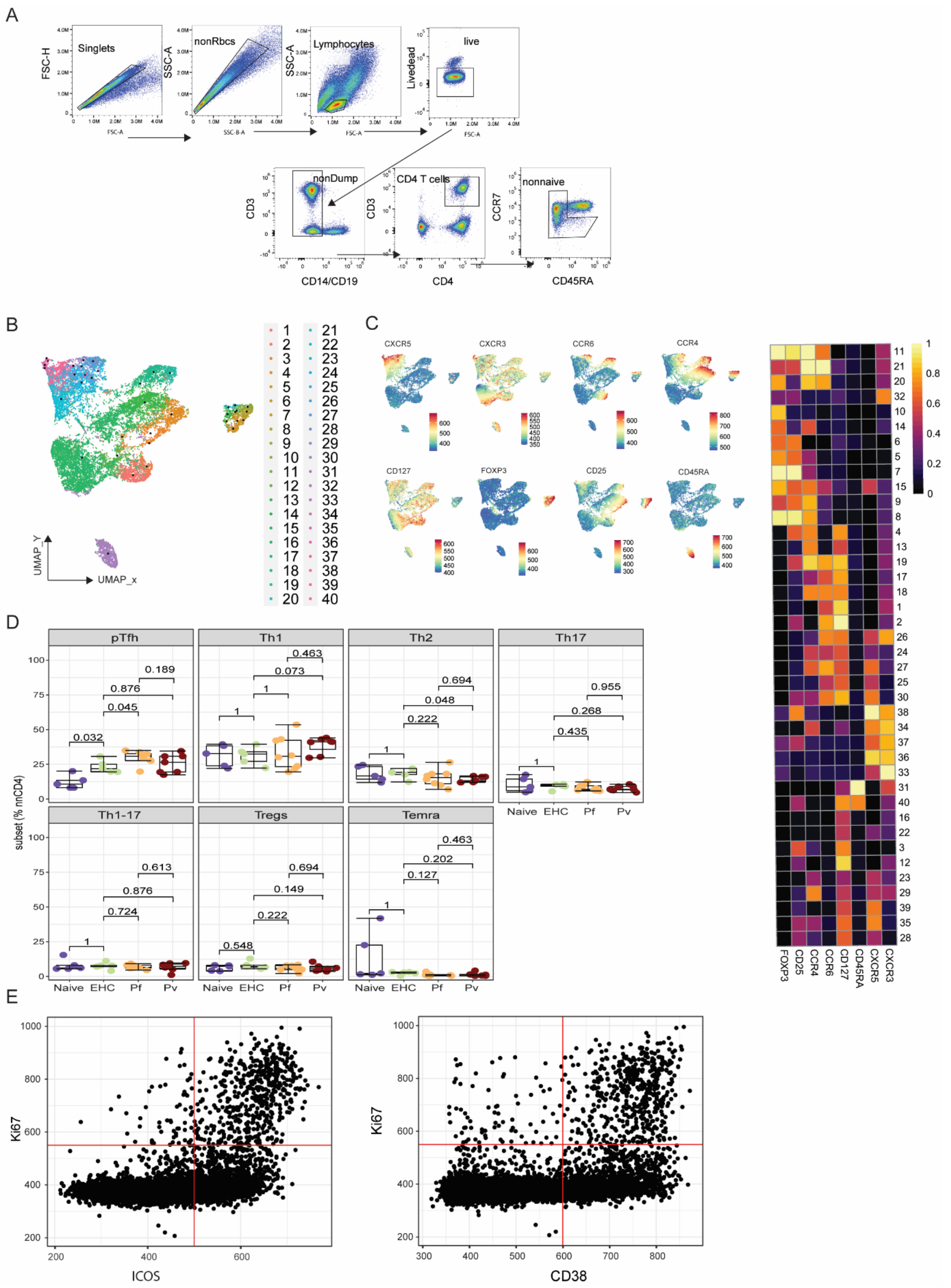


***Supplementary Figure S3: Activation of CD4 T cell subsets (Sub-Cohort 2)***

*CD4 T cells were analysed with a comprehensive panel and unbiased approaches in healthy malaria naïve Australians, healthy malaria uninfected controls, P. falciparum and P. vivax patients (Naïve, n=5; EC, n=5; P. falciparum (Pf), n = 8; P. vivax (Pv), n = 7).* ***A)*** *Gating* *strategy for identification of nonnative CD4 T cells.* ***B)*** *Unsupervised clustering with cell clusters visualised on a UMAP.* ***C)*** *Expression of markers used to annotate clusters shown on a UMAP and heatmap.* ***D)****Frequencies of subsets as proportions of CD4 T cells.* ***E)*** *Gating strategy to identify ICOS+, CD38+ and Ki67+ cells in Spectre. Data in box plots are individual responses coloured according to the control and malaria groups. Box and whisker plots indicate first and third quartiles for the hinges, median line, and lowest and highest values no further than 1.5 interquartile range from the hinges for whisker lines. P-values are Wilcoxon signed-rank test*

#### Supplementary Tables and Figures

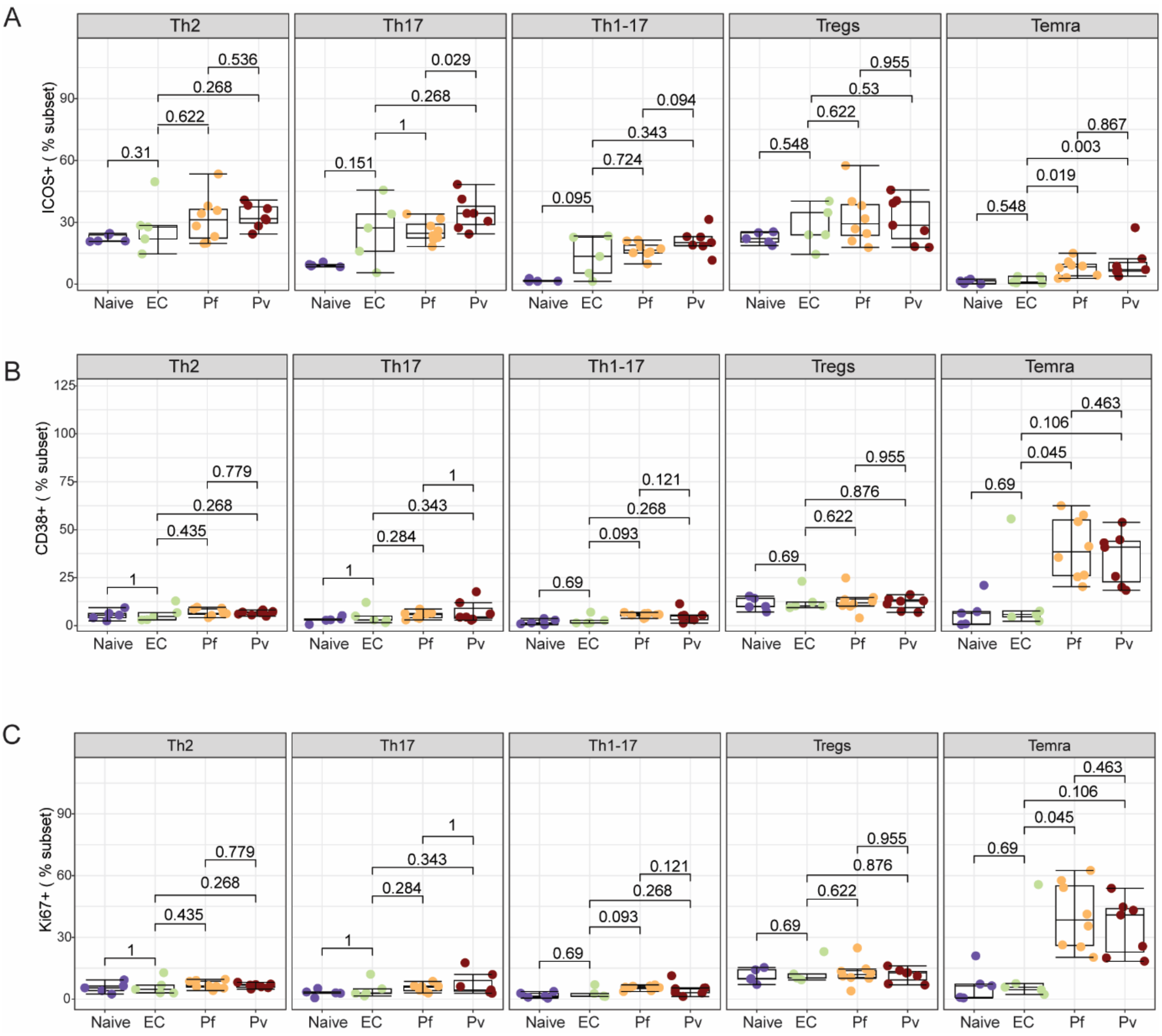


***Supplementary Figure S4: Activation of CD4 T cell subsets (Sub-Cohort 2)***

*CD4 T cells were analysed with a comprehensive panel and unbiased approaches in healthy malaria naïve Australians, healthy malaria uninfected controls, P. falciparum and P. vivax patients (Naïve, n=5; EC, n=5; P. falciparum (Pf), n = 8; P. vivax (Pv), n = 7). Frequency of activated A**) ICOS+,* ***B)*** *CD38+, and* ***C)*** *Ki67+ cells as proportions of CD4 T cell subsets. Data are individual responses coloured according to the control and malaria groups. Box and whisker plots indicate first and third quartiles for the hinges, median line, and lowest and highest values no further than 1.5 interquartile range from the hinges for whisker lines. P-values are Wilcoxon signed-rank test.*

#### Supplementary Tables and Figures

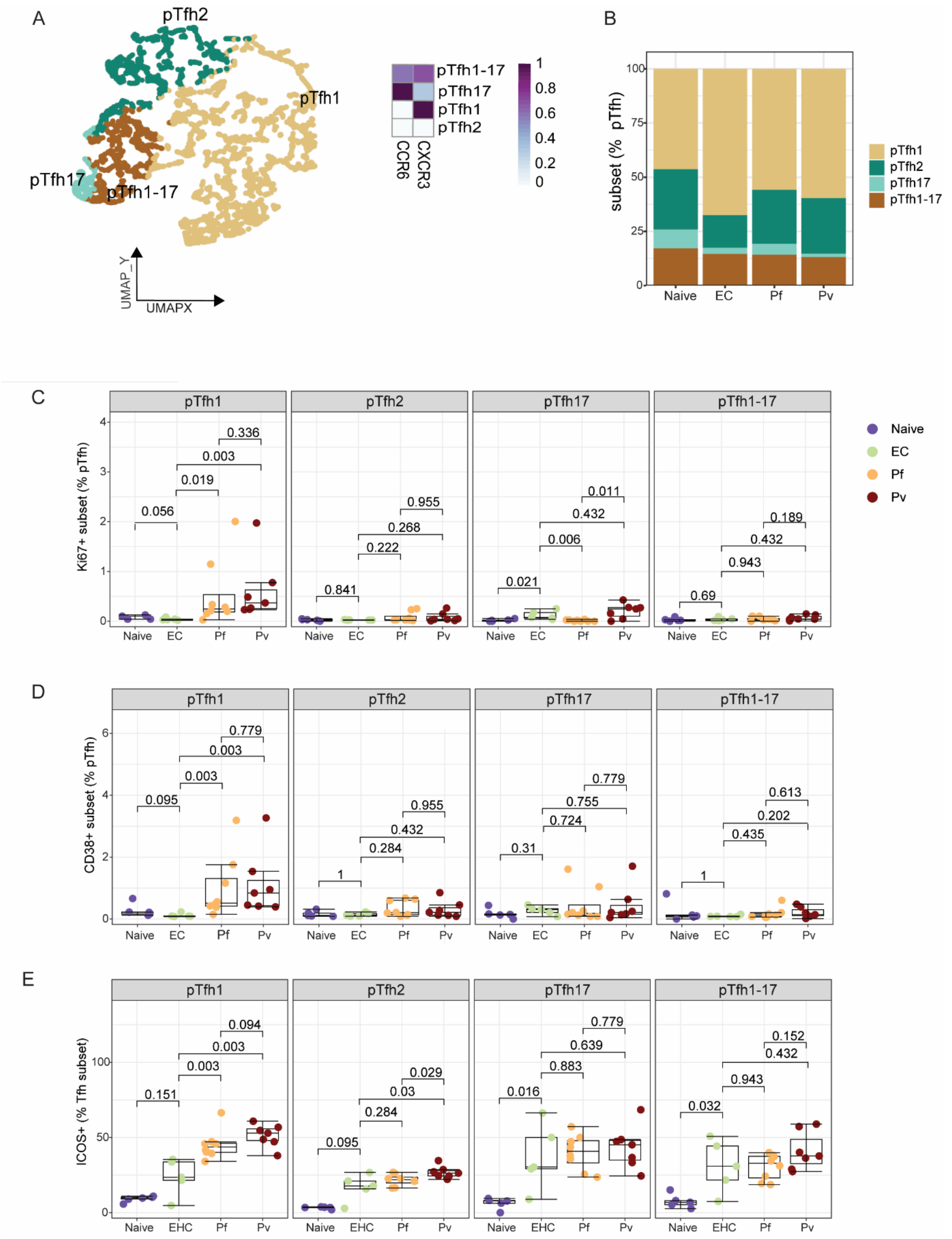


***Supplementary Figure S5: pTfh gating and subset activation (Sub-Cohort 2)***

*pTfh cell subsets were identified by unsupervised clustering assessed in adults with P. falciparum (Pf, n = 25) and P. vivax (Pv, n = 12) compared with endemic healthy controls (EC, n = 8).* ***A)*** UMAP showing clusters of pTfh cell subsets based on the expression of markers in the heatmap. ***B)*** *Ki67+ cells as proportions of total pTfh in controls and malaria* *patients.* ***C)*** *ICOS+ and* ***D)*** *Ki67+ cells within pTfh subsets. For all plots, box and whisker plots indicate first and third quartiles for the hinges, median line, and lowest and highest values no further than 1.5 interquartile range from the hinges for whisker lines. P-values are Wilcoxon signed-rank test.*

**Supplementary Tables and Figures**

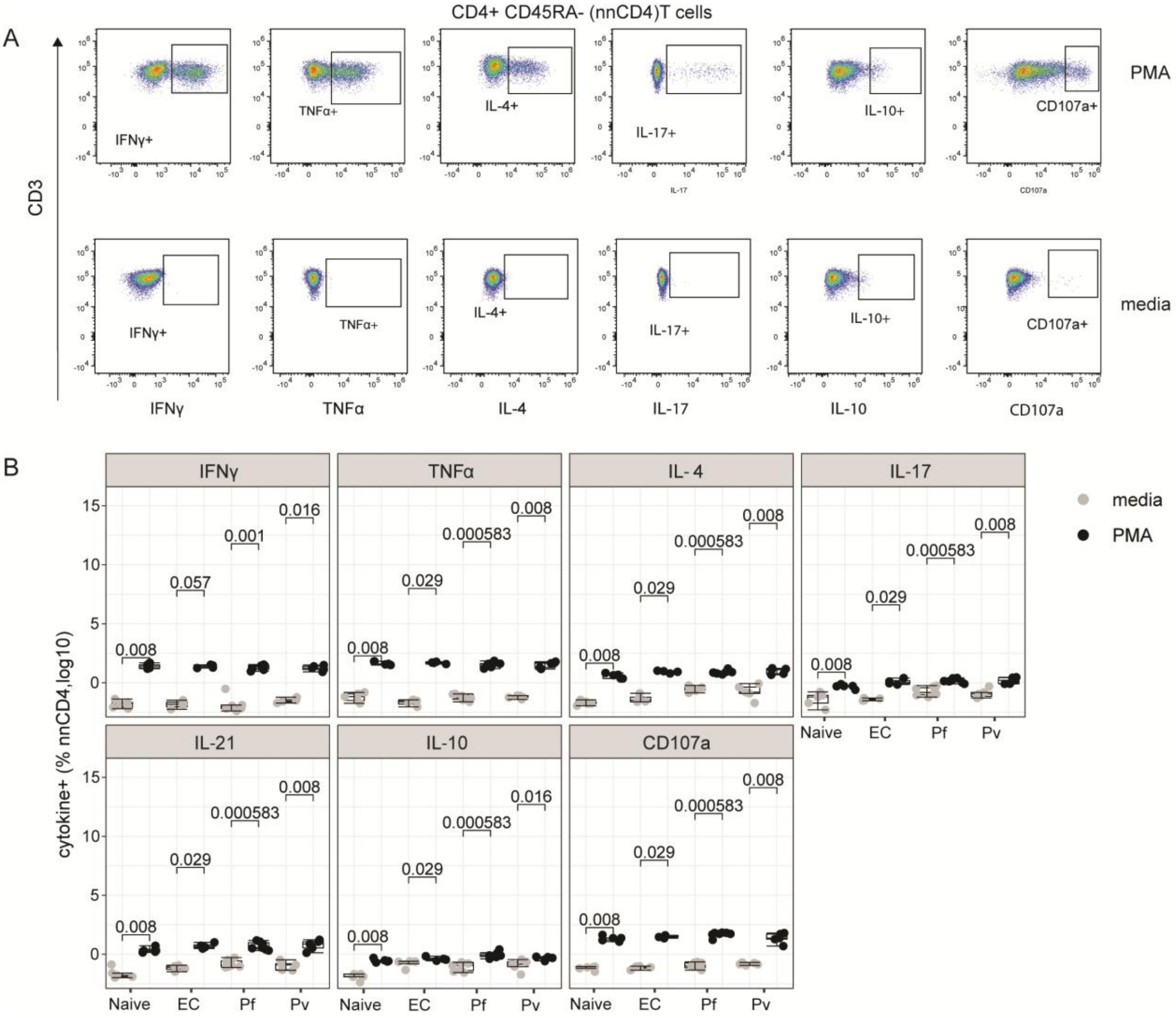


***Supplementary Figure: 6 Gating for PMA cytokine staining and CD4 T cell cytokine expression***

*PBMC from healthy malaria naïve Australians (Naïve, n = 5), endemic healthy controls (EHC, n = 5), P. falciparum (Pf, n = 8) and P. vivax (Pv, n = 7)infected adults were stimulated with PMA/Ionomycin(PMA/I) or unstimulated (media) and IFNγ, TNFα, IL-4, IL-17, IL-21, IL-10 production and CD107a expression secretion determined by intracellular staining.* ***A)*** *Gating strategy to identify cytokine expression.* ***B)*** *Comparison of the log10 frequency of IFNγ, TNFα, IL-4, IL-17, IL-21, IL-10 producing and CD107a expressing nnCD4 T cells between PMA stimulated (black) and no stimulation (grey) in controls and patients. Box and whisker plots indicate first and third quartiles for the hinges, median line, and lowest and highest values no further than 1.5 interquartile range from the hinges for whisker lines. P-values are Wilcoxon signed-rank test*

#### Supplementary Tables and Figures

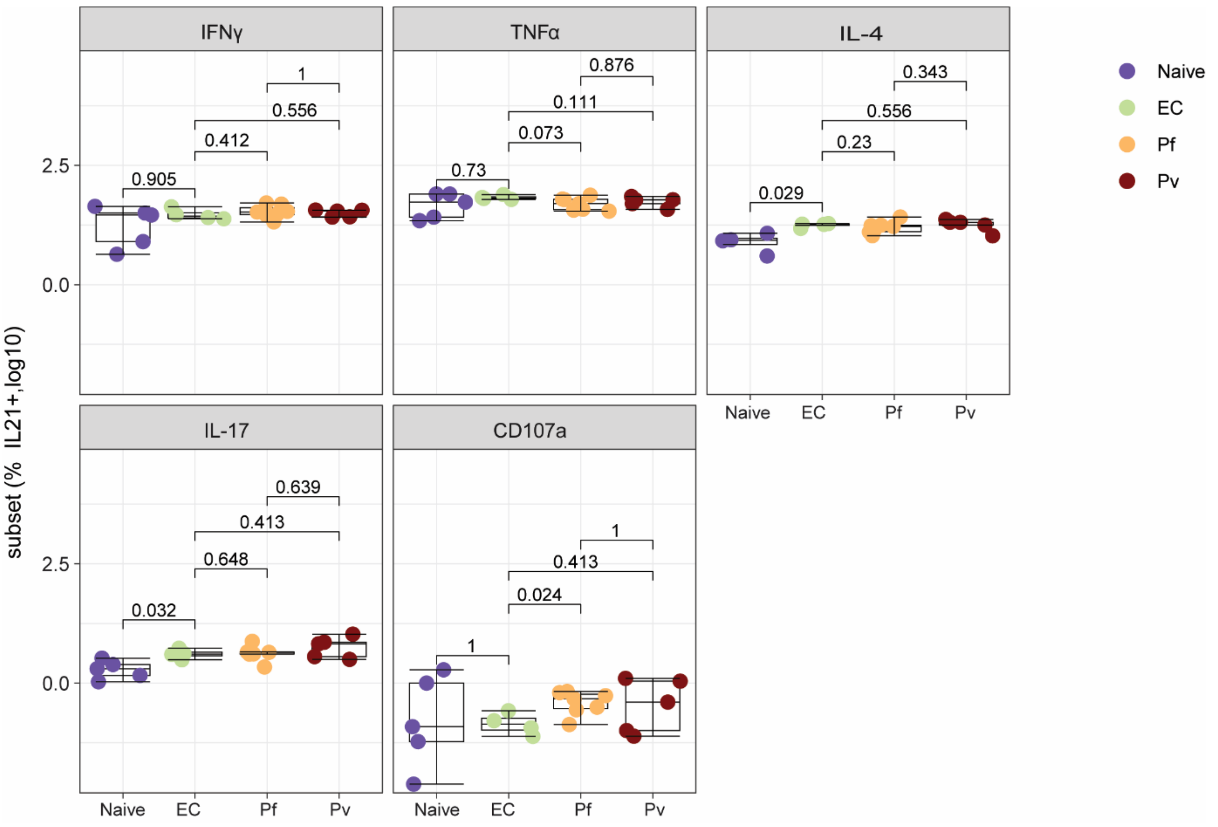


***Supplementary Figure 7: Co-expression of IL21 with other cytokines***

*Comparison of the log10 frequency of IL-21+ CD4 T cell co-production with IFNγ, TNFα, IL-4, IL-17, IL-21, IL-10  and CD107a expression following stimulation with PMA/Io in malaria naive Australians (Naïve, n = 5), endemic healthy controls (EC, n = 5), P. falciparum (Pf, n = 8) and P. vivax (Pv, n = 7)infected adults.*
